# The Omics Discovery REST interface

**DOI:** 10.1101/2020.02.10.939967

**Authors:** Gaurhari Dass, Manh-Tu Vu, Pan Xu, Enrique Audain, Marc-Phillip Hitz, Henning Hermjakob, Yasset Perez-Riverol

**Affiliations:** European Molecular Biology Laboratory, EMBL-European Bioinformatics Institute (EMBL-EBI), Cambridge CB10 1SD, UK; State Key Laboratory of Proteomics, Beijing Proteome Research Center, Beijing Institute of Lifeomics, National Center for Protein Sciences (The PHOENIX Center, Beijing), 102206, Beijing, China; Department of Human Genetics, University Medical Center Schleswig-Holstein (UKSH), Kiel, Germany

**Keywords:** multiomics, omics data, FAIR principles, REST, application programming interface (API)

## Abstract

The Omics Discovery Index is an open source platform that can be used to access, discover and disseminate omics datasets. OmicsDI integrates proteomics, genomics, metabolomics, models and transcriptomics datasets. Using an efficient indexing system, OmicsDI integrates different biological entities including genes, transcripts, proteins, metabolites and the corresponding publications from PubMed. In addition, it implements a group of pipelines to estimate the impact of each dataset by tracing the number of citations, reanalysis and biological entities reported by each dataset. Here, we present the OmicsDI REST interface to enable programmatic access to any dataset in OmicsDI or all the datasets for a specific provider (database). Clients can perform queries on the API using different metadata information such as sample details (species, tissues, etc), instrumentation (mass spectrometer, sequencer), keywords and other provided annotations. In addition, we present two different libraries in R and Python to facilitate the development of tools that can programmatically interact with the OmicsDI REST interface.

## Introduction

In recent years, major advances in the field of omics analyses have led to an exponential increase in available experimental data (1). Omics platforms offer high-throughput, detailed exploration of the genome, transcriptome, proteome and metabolome, analysed using a variety of techniques including mRNA and miRNA arrays, next-generation sequencing and mass spectrometry (2,3). The development of new analytical methods and instruments has enabled the analysis of one biological sample to generate many kinds of “big” omics data in parallel, such as genome sequence (genomics), patterns of gene and protein expression (transcriptomics and proteomics), and metabolite concentrations (metabolomics) (4). In addition, public data deposition is growing in all omics disciplines, because it is considered to be a good scientific practice (e.g. to enable reproducibility) and/or it is mandated by funding agencies and scientific journals (5). These new developments haves triggered new challenges and opportunities to make data Findable, Accessible, Interoperable and Re-usable (FAIR - https://www.force11.org/group/fairgroup/fairprinciples) (6).

In 2016, we released the first version of the Omics Discovery Index (OmicsDI— https://www.omicsdi.org) as a light-weight system to aggregate datasets across multiple public omics data resources (1,5). OmicsDI collects datasets from multiple repositories and databases representing genomics, transcriptomics, proteomics, metabolomics and multiomics datasets, as well as computational models of biological processes. Datasets can be searched and filtered based on different types of technical and biological annotations (e.g. species, tissues, diseases, etc.), year of publication and the original data repository where they are stored, among others. As of January 2020, OmicsDI stores just over 453,900 datasets from 20 different public data resources (https://www.omicsdi.org/database). The OmicsDI web interface provides different views and search capabilities on the indexed datasets. In addition, every dataset is encoded in the web interface using schema.org (https://schema.org/) representation for datasets enabling resources such as Google datasets (https://datasetsearch.research.google.com/) to index omics data.

The FAIR principles are aimed at computational interfaces as well as web interfaces for human use. In this manuscript we describe the main features of the Omics Discovery Index interface (API), which enables programmatic access to OmicsDI datasets (https://www.omicsdi.org/ws). The API allows programmatic access to all OmicsDI datasets and the corresponding information including metadata, data files and molecules reported by the datasets. In addition, it allows to perform search and filters on the datasets by different properties such as tissue, cell type, or organisms. This API is used by different external resources, such as DataMed (https://datamed.org/) (7) or MENDA (http://menda.cqmu.edu.cn:8080/index.php) (8). Finally, we introduce two new client libraries in R (ddiR - https://github.com/OmicsDI/ddiR) and python (ddipy - https://github.com/OmicsDI/ddipy) to enable bioinformaticians and developers to develop new tools and packages that interact with OmicsDI.

## Design and Implementation

The OmicsDI API is implemented in Java, building on top of the Spring framework (http://projects.spring.io/spring-framework/). Data queries are powered by optimized Apache Solr servers (http://lucene.apache.org/solr/). Data can be accessed over HTTP (HyperText Transfer Protocol) via REST-like ‘Get’ requests, which ensures that the services are easy to use and are supported by all major platforms. JSON (JavaScript Object Notation) and XML (Extensible Markup Language) were chosen as the output format of a common metadata representation for all omics types (e.g. proteomics, genomics or transcriptomics) (1).

All OmicsDI API (https://www.omicsdi.org/ws) methods can be classified into five major categories (**Table 1**); including datasets, database, and statistics. Table 1 summarises the most relevant and widely used methods from the API organized by categories.

**Table 1:** Most relevant methods provided by OmicsDI REST Interface.

| *Category* | *Method* | *Description* | *Type of Method* |
| --- | --- | --- | --- |
| *Dataset* | /dataset/search | Search for datasets in OmicsDI | GET |
|  | /dataset/{database}/{accession} | Retrieve a specific dataset from OmicsDI | GET |
|  | /dataset/batch | Retrieve a batch of datasets | GET |
|  | /dataset/getFileLinks | Retrieve all file links for a given dataset | GET |
|  | /dataset/{database}/{accession}/files | Retrieve the list of dataset's files | GET |
|  | /dataset/merge | Merge two different replicate datasets | POST |
|  | /dataset/getMergeCandidates | Get all the datasets that can be consider as replicates | GET |
|  | /dataset/latest | Retrieve the latest datasets added to OmicsDI | GET |
|  | /dataset/getSimilarByPubmed | Retrieve similar datasets based on PubMed identifier | GET |
|  | /dataset/getSimilar | Retrieve the related datasets to one Dataset | GET |
| *Database* | /database/all | Retrieve OmicsDI databases/repositories | GET |
| *Ontology Terms* | /term/getTermByPattern | Search dictionary Terms | GET |
|  | /term/frequentlyTerm/list | Retrieve frequently terms | GET |
| *Statistics* | /statistics/organisms | Return statistics per organisms | GET |
|  | /statistics/tissues | Return statistics per tissues | GET |
|  | /statistics/omics | Return statistics per Omics type | GET |
|  | /statistics/diseases | Return statistics per diseases | GET |
|  | /statistics/domains | Return statistics per Repository | GET |
| *SEO (Schema.org)* | /seo/home | Retrieve JSON+LD for home page | GET |
|  | /seo/search | Retrieve JSON+LD for browse page | GET |
|  | /seo/api | Retrieve JSON+LD for api page | GET |
|  | /seo/schema/{database}/{accession} | Retrieve JSON+LD Schema for dataset page | GET |
|  | /seo/database | Retrieve JSON+LD for databases page | GET |
|  | /seo/dataset/{database}/{accession} | Retrieve JSON+LD for dataset page | GET |
|  | /seo/about | Retrieve JSON+LD for about page | GET |

The *datasets* category includes all the methods to enable search and retrieval of the datasets from OmicsDI. For example, the most extensively use method in the API (*/dataset/search*) allows users to search datasets by different attributes including tissues, organism or instrument. The search entry point *(/dataset/search*) uses different parameters to sort, query, facet count, and paginate through the results.

The query value is used to query all datasets that contain the specific keyword. For example, https://www.omicsdi.org/ws/dataset/search?query=human will retrieve all the datasets that contain the word “human” in any part of the metadata. Multiple search terms separated by white spaces are combined by default in AND logic. Therefore, an input text containing for example glutathione transferase is treated as glutathione AND transferase and only entries having both terms will be found (see query language syntax - https://www.ebi.ac.uk/ebisearch/documentation.ebi#query_syntax). It is important to notice that the default order of results is based on their relevance. Then, the API will return first all the datasets that contains both words and then the datasets that contains at least one of them.

If the query keyword is to be applied to one specified field, the keyword should be combined with the field. For example, if the user wants to query all the datasets where the protein has been identified, the uniport accession should be combined with the field UNIPROT like: https://www.omicsdi.org/ws/dataset/search?query=UNIPROT:P21399.

The method */dataset/getSimilar* retrieve all the datasets that are similar to a specific dataset by metadata (see original manuscript of OmicsDI for the similarity algorithm explanation (1)). Another important method */dataset/getMergeCandidates* enables to retrieve candidate datasets that can be considered as replicates (5).

The *seo* category enables to retrieve all the information from datasets and databases using *schema.org* representation. These methods are designed for future resources and services that use schema.org and BioSchemas (https://bioschemas.org/) for crawling and indexing. While currently Google Datasets (https://datasetsearch.research.google.com/) is already crawling all OmicsDI datasets using the web application, future services may use the OmicsDI Rest interface for large scale indexing rather than web crawling.

All the API methods returning lists of objects can be navigated using pagination with two parameters: *start* – number of the page, and *size* – number of objects to retrieve in the page.

## Data Files Geolocation

OmicsDI Rest API */dataset/{database}/{accession}* method allows to get the information for one particular dataset including the metadata and public URI (Uniform Resource Identifier) of the data files. An important feature of this method is to be able to provide the URI of the files that are closer to the user query (**Figure 1**). OmicsDI stores for each dataset a primary source of the resource and all the replicates of it to avoid duplications when the dataset is replicated in multiple providers, for example, GEO and ArrayExpress, or PRIDE and MassIVE (5).

**Figure 1:**
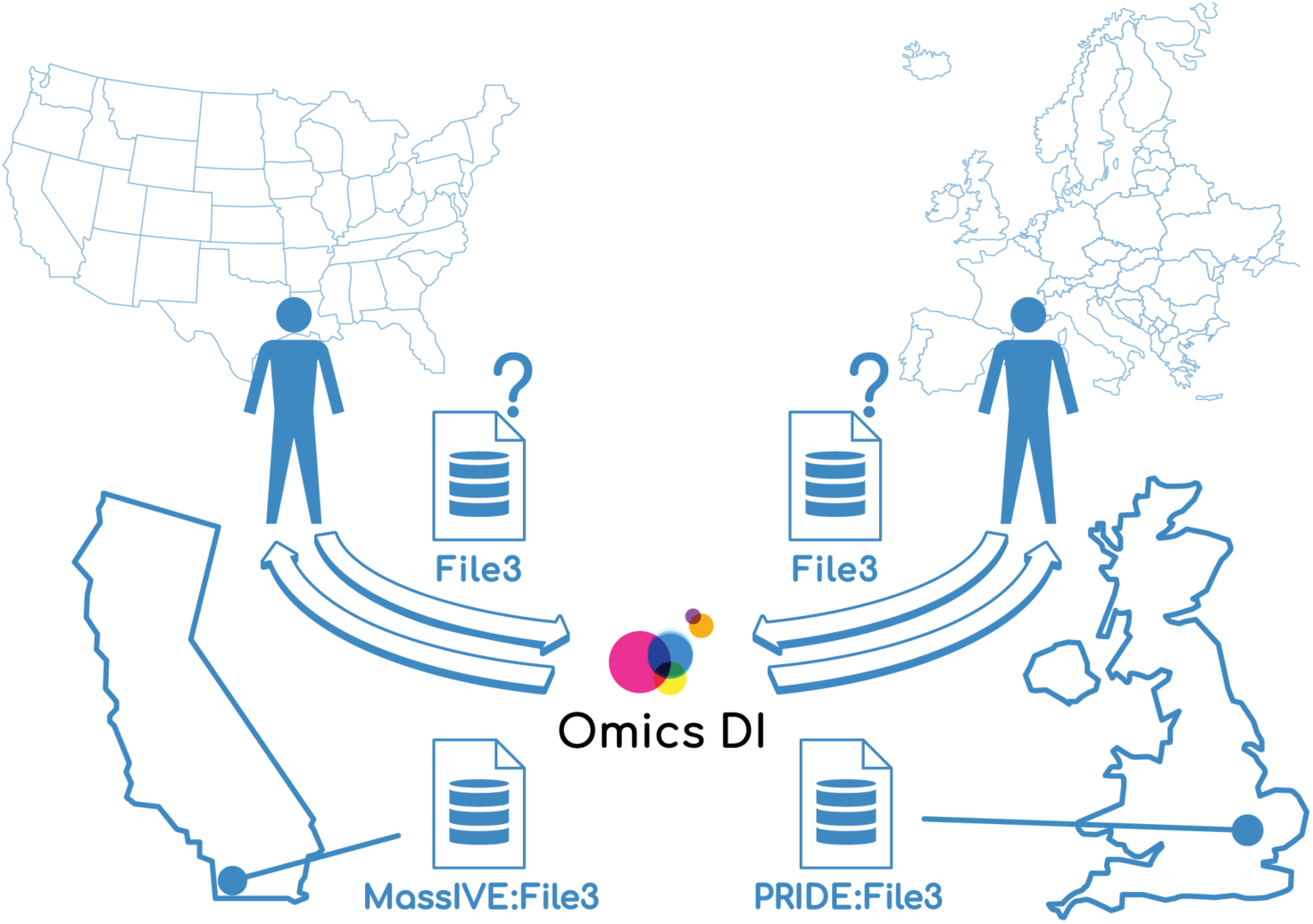
Data file geolocation feature in OmicsDI Rest API. The user request localization define the files or copy of the dataset that will be retrieve by the API.

When two versions of the same dataset are available in two different databases, the API highlights the files that are closer to the client request as primary. The method */dataset/{database}/{accession}* uses the attribute from the header request “X-Forwarded-For: IP”, to compute how close the request is to some of the providers. This method enables the user to download the data files closer to them. For example, if the request is performed from the United States, the replicate from MassIVE will be marked as primary and the PRIDE files as mirror, if the request is performed from Germany the PRIDE datasets will be marked as primary and MassIVE as mirror (**Figure 1**).

## Example use cases

A user of the web services might be interested in retrieving multiomics projects that contain proteomics and transcriptomics data. Since the omics_type stores the omics type of the dataset, the user can perform the following query:

https://www.omicsdi.org/ws/dataset/search?query=omics_type:%22Transcriptomics%22%20AND%20omics_type:%22Proteomics%22. By combining two different values for the same field omics_type the user can query all the datasets that contains both values transcriptomics and proteomics. The user can use all the fields defined by the OmicsDI dataset schema (dataset schema - https://github.com/OmicsDI/specifications/blob/master/docs/schema/fields.md) to refine their filters.

Examining the details of one specified dataset from the results query can be done by calling the dataset method, for example https://www.omicsdi.org/ws/dataset/arrayexpress-repository/E-GEOD-53085. To then retrieve some specific files from the dataset, the user could use the method https://www.omicsdi.org/ws/dataset/arrayexpress-repository/E-GEOD-53085/files?position=1,2,3,4,5,6. This method (dataset/{database}/{accession}/files) uses the parameter position to enable filtering the files related with the project by file index.

Another interesting use case is finding all the datasets where a specific molecule, such as a protein, gene or metabolite, has been reported as identification or differentially expressed in a dataset. For example, a user can be interested in all the proteomics projects where the UniProt proteins “Q99714” and “P06744” have been identified: https://www.omicsdi.org/ws/dataset/search?query=UNIPROT:P06744%20AND%20UNIPROT:Q99714%20AND%20omics_type:Proteomics.

The same type of queries can be performed for other omics types. For example, if the user is interested in all the transcriptomics experiments where gene ENSG00000147251 significantly changes expression, the following query can be used: https://www.omicsdi.org/ws/dataset/search?query=ENSEMBL:ENSG00000147251.

As discussed above, one of the interesting features of OmicsDI is the possibility to retrieve the information about replicates of the same dataset (5). For example, the dataset https://www.omicsdi.org/dataset/pride/PXD003213 is stored in two different places: PRIDE (9) and MassIVE (10). The OmicsDI REST interface merges all the replicates of a dataset in the same dataset entry but allows the users to download the files from the replicate that is closer by region to the client request. Figure 2 shows two different json responses depending on the location of the request. When the query is performed from an IP in United States the primary files are the ones from MassIVE (**Figure 2a**); in contrast; when the query is performed from an IP in United Kingdom the primary files are the ones from PRIDE (**Figure 2b**).

**Figure 2:**
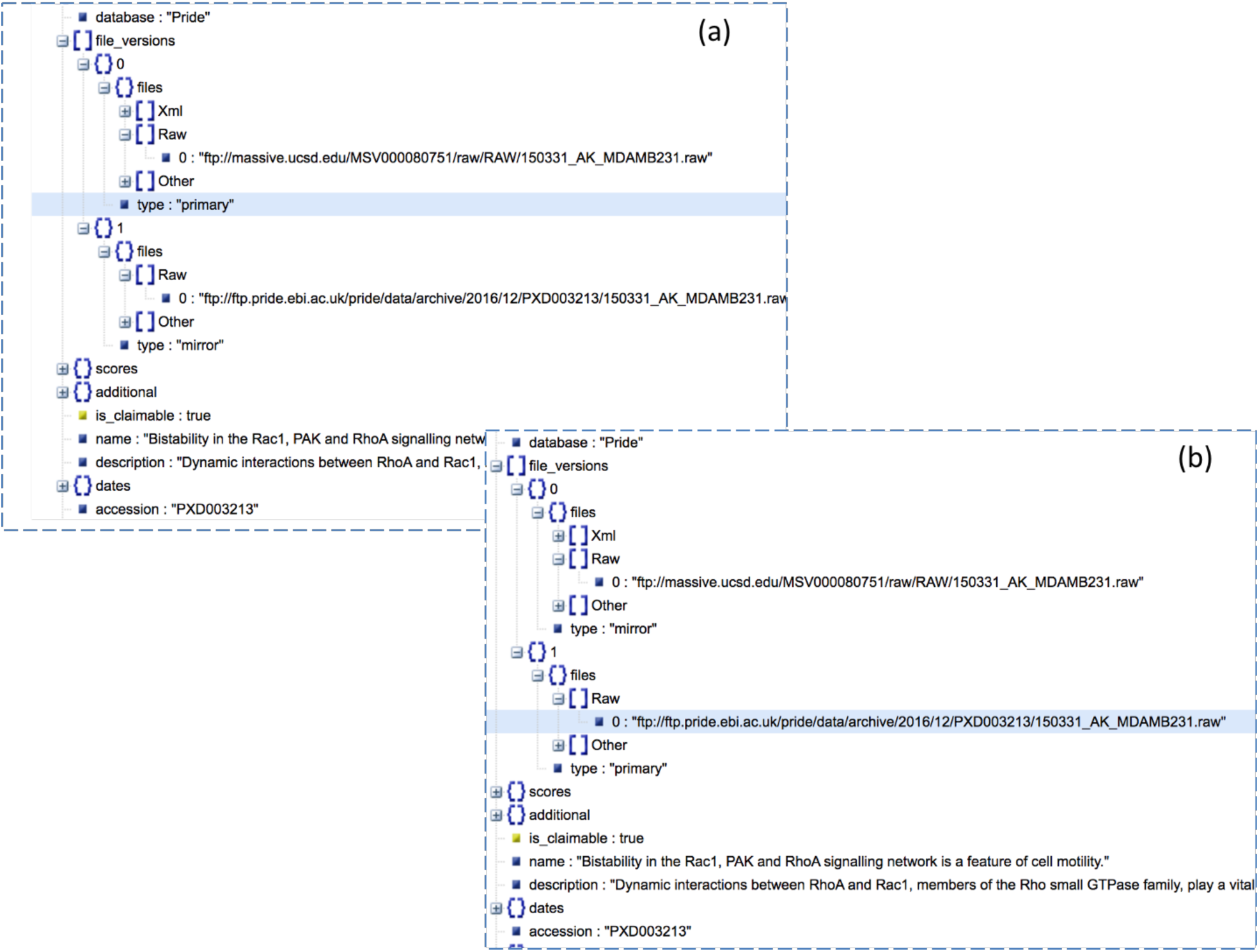
The OmicsDI Rest Interface enables to retrieve from each dataset the files closest to the IP client request. For example, for dataset https://www.omicsdi.org/dataset/pride/PXD003213 the API marks as primary source the (a) MassIVE files when the query is performed from United States (*curl --header “X-Forwarded-For: 66.165.239.58” https://www.omicsdi.org/ws/dataset/pride/PXD003213?);* and (b) PRIDE files are marked as primary when the request is performed from United Kingdom *(curl --header “X-Forwarded-For: 193.62.193.80” https://www.omicsdi.org/ws/dataset/pride/PXD003213*).

## The OmicsDI REST clients

In order to facilitate the development of new tools and services that use the OmicsDI Rest interface, we have implemented two libraries in R (ddiR - https://github.com/OmicsDI/ddiR) and python (ddipy - https://github.com/OmicsDI/ddipy). Both libraries provide methods and data structures to interact with the Rest API; including methods to search, navigate and retrieve datasets and files. For example, if the developer would like to retrieve and specific dataset information from the API:

In python:

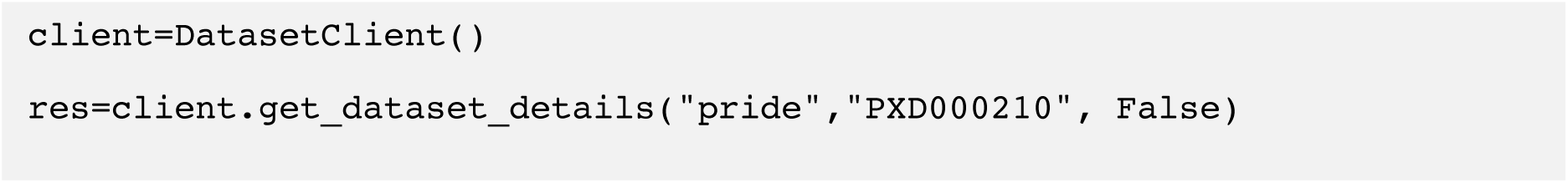

In R:

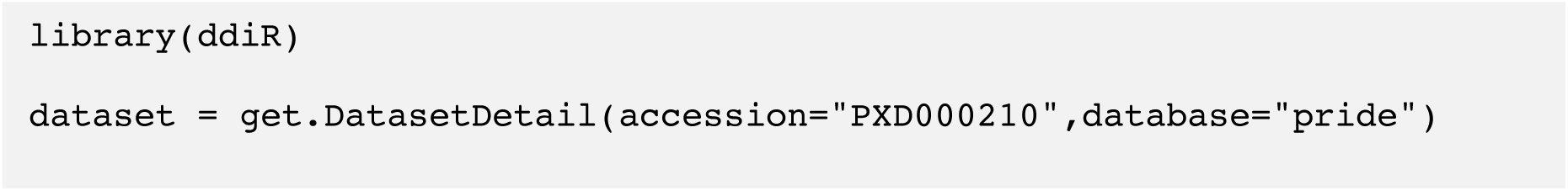

The R package is implemented using S4 object orientation, and the methods are documented using roxygen comments. The R package can be installed using devtools like *install_github(“OmicsDI/ddiR”)*. The python package has been added to pip and can be installed using *pip install ddipy*.

## Discussion

The OmicsDI REST interface has been developed to enable programmatic access to the OmicsDI database and all the datasets provided by OmicsDI partners and repositories. With the increase of the number of these resources and the creation of new national infrastructures for personal genomics data storage and dissemination, a central system like OmicsDI that indexes the metadata can help users to find and retrieve distributed datasets (11-13). Currently, OmicsDI is working on implementing the GA4GH (Global Alliance for Genomics and Health) repository service specification (https://github.com/ga4gh/data-repository-service-schemas) to enable genomics workflows and cloud infrastructures to retrieve the datasets closest to the compute platform using the current geolocation data file feature.

The API has been available since April 2017 and a few internal and external applications are already making use of the new functionality: DataMed (https://datamed.org/) (7) or MENDA (http://menda.cqmu.edu.cn:8080/index.php) (8). Since the first release, the OmicsDI datasets has have been visited in the OmicsDI web site 1,233,388, 253,428, 435,859, 860,092, 1,417,107, 14,793,937 times for genomics, metabolomics, models, multiomics, proteomics and transcriptomics, respectively. With the new R and python libraries, we facilitate the development of new tools and packages that can use the OmicsDI REST interface to search and retrieve datasets from OmicsDI. The OmicsDI REST Interface will continue to develop in parallel with the OmicsDI web page. Should users wish to discuss requests for new functionality, the authors encourage them to contact the OmicsDI helpdesk with their suggestions.

## Acknowledgments

G.D. is supported by EMBL core funding. P.X. was supported by BBSRC International Partnering Award BB/N022432/1. Y.P.-R. acknowledges the Wellcome Trust (grant number 208391/Z/17/Z) and the EPIC-XS project (grant number 823839), funded by the Horizon 2020 programme of the European Union.

## References

1. Perez-Riverol, Y., Bai, M., da Veiga Leprevost, F., Squizzato, S., Park, Y.M., Haug, K., Carroll, A.J., Spalding, D., Paschall, J., Wang, M. et al. (2017) Discovering and linking public omics data sets using the Omics Discovery Index. Nat Biotechnol, 35, 406–409.

2. Perez-Riverol, Y., Kuhn, M., Vizcaino, J.A., Hitz, M.P. and Audain, E. (2017) Accurate and fast feature selection workflow for high-dimensional omics data. PLoS One, 12, e0189875.

3. Manzoni, C., Kia, D.A., Vandrovcova, J., Hardy, J., Wood, N.W., Lewis, P.A. and Ferrari, R. (2018) Genome, transcriptome and proteome: the rise of omics data and their integration in biomedical sciences. Brief Bioinform, 19, 286–302.

4. Field, D., Sansone, S.A., Collis, A., Booth, T., Dukes, P., Gregurick, S.K., Kennedy, K., Kolar, P., Kolker, E., Maxon, M. et al. (2009) Megascience. ‘Omics data sharing. Science, 326, 234–236.

5. Perez-Riverol, Y., Zorin, A., Dass, G., Vu, M.T., Xu, P., Glont, M., Vizcaino, J.A., Jarnuczak, A.F., Petryszak, R., Ping, P. et al. (2019) Quantifying the impact of public omics data. Nat Commun, 10, 3512.

6. Wilkinson, M.D., Dumontier, M., Aalbersberg, I.J., Appleton, G., Axton, M., Baak, A., Blomberg, N., Boiten, J.W., da Silva Santos, L.B., Bourne, P.E. et al. (2016) The FAIR Guiding Principles for scientific data management and stewardship. Sci Data, 3, 160018.

7. Ohno-Machado, L., Sansone, S.A., Alter, G., Fore, I., Grethe, J., Xu, H., Gonzalez-Beltran, A., Rocca-Serra, P., Gururaj, A.E., Bell, E. et al. (2017) Finding useful data across multiple biomedical data repositories using DataMed. Nat Genet, 49, 816–819.

8. Pu, J., Yu, Y., Liu, Y., Tian, L., Gui, S., Zhong, X., Fan, C., Xu, S., Song, X., Liu, L. et al. (2019) MENDA: a comprehensive curated resource of metabolic characterization in depression. Brief Bioinform.

9. Perez-Riverol, Y., Csordas, A., Bai, J., Bernal-Llinares, M., Hewapathirana, S., Kundu, D.J., Inuganti, A., Griss, J., Mayer, G., Eisenacher, M. et al. (2019) The PRIDE database and related tools and resources in 2019: improving support for quantification data. Nucleic Acids Res, 47, D442–D450.

10. Wang, M., Carver, J.J., Phelan, V.V., Sanchez, L.M., Garg, N., Peng, Y., Nguyen, D.D., Watrous, J., Kapono, C.A., Luzzatto-Knaan, T. et al. (2016) Sharing and community curation of mass spectrometry data with Global Natural Products Social Molecular Networking. Nat Biotechnol, 34, 828–837.

11. Siu, L.L., Lawler, M., Haussler, D., Knoppers, B.M., Lewin, J., Vis, D.J., Liao, R.G., Andre, F., Banks, I., Barrett, J.C. et al. (2016) Facilitating a culture of responsible and effective sharing of cancer genome data. Nat Med, 22, 464–471.

12. Chalmers, D.R., Nicol, D. and Otlowski, M.F. (2014) To share or not to share is the question. Appl Transl Genom, 3, 116–119.

13. Fiume, M., Cupak, M., Keenan, S., Rambla, J., de la Torre, S., Dyke, S.O.M., Brookes, A.J., Carey, K., Lloyd, D., Goodhand, P. et al. (2019) Federated discovery and sharing of genomic data using Beacons. Nat Biotechnol, 37, 220–224.

